# Mono-Association with *Lactobacillus plantarum* Disrupts Intestinal Homeostasis in adult *Drosophila*

**DOI:** 10.1101/049981

**Authors:** David Fast, Aashna Duggal, Edan Foley

## Abstract

The microbiome of *Drosophila* promotes intestinal stem cell division through evolutionarily conserved biochemical pathways. As such, axenic flies have lower rates of gut stem cell division than age-matched wild type counterparts. Additionally, flies with a full consortium of symbiotic bacteria are shorter lived than those maintained in the absence of a microbiome. However, we do not know if stem cell division is essential for symbiont-dependent regulation of adult fly lifespan. To determine if individual symbionts cause aging-dependent death in *Drosophila*, we examined the impacts of common symbionts on host longevity. In this study, we found that mono-association of adult *Drosophila* with *Lactobacillus plantarum*, a widely reported fly symbiont, and member of the probiotic *Lactobacillus* genus, curtails adult longevity relative to germ-free counterparts. However, the effects of *plantarum* on lifespan were independent of intestinal aging. Instead, we found that association with *plantarum* causes an extensive intestinal pathology within the host, characterized by loss of intestinal stem cells, impaired epithelial renewal, and a gradual erosion of epithelial integrity. Our study uncovers an unknown aspect of *Lactobacillus plantarum*-*Drosophila* interactions, and establishes a simple model to characterize symbiont-dependent disruption of intestinal homeostasis.

## INTRODUCTION

Environmental, microbial, and host factors establish an intestinal environment that permits gut colonization by a variable consortium of bacteria. Extrinsic factors such as pH, oxygen, and nutrient supply influence the biogeography of microbe distribution, physical barriers contain microbes to the gut lumen, and bacteriostatic products such as antimicrobial peptides and reactive oxygen species block bacterial dissemination throughout the host. Inside the lumen, microbes compete with each other for access to nutrients and intestinal attachments sites, and release metabolites that influence processes as diverse as growth, immunity, and behavior in the host (1–6). Shifts in the composition or distribution of the bacterial community often lead to the onset of debilitating, and potentially deadly, diseases for the hosts (4, 7, 8).

*Drosophila melanogaster* is a useful model for studies that examine direct interactions between a host and individual symbiotic species (9–13). The fly microbiome consists of a limited number of bacterial species that are easily cultured and manipulated in isolation (14–19). Researchers have access to simple protocols for the establishment of gnotobiotic fly cultures, and flies lend themselves to sophisticated manipulation of host gene expression (13, 20). Of equal importance, there are extensive genetic, developmental, and biochemical similarities between fly and mammalian gut biology (21–23). Thus, discoveries in *Drosophila* provide insights into evolutionarily conserved features of host-bacteria interactions. For example, in flies and mammals, basal Intestinal Stem Cells (ISCs) divide and differentiate at a rate that maintains an intact epithelial barrier (20, 22, 24–26). A relatively simple “escalator” program matches ISC division to loss of aged cells, while a more complex, adaptive program activates ISC division to compensate for environmental destruction of host cells (27). This adaptive regulation of growth maintains the integrity of the epithelial barrier, and is critical for long-term health of the host. Breaches to the gut barrier permit an invasion of intestinal microbes that activate local immune responses, and drive the development of chronic inflammatory illnesses (4, 28–30)

Although the microbiome of *Drosophila* is orders of magnitudes less complex than that found in mammals (14, 18) populations of *Lactobacillus* species are common to fly midguts and animal small intestines (14, 18, 20). Studies of the *Drosophila* symbiont, *Lactobacillus plantarum* (*Lp*), uncovered several interactions between the two species. *Lp* contributes to larval growth (9), uptake of dietary protein (31), and management of malnutrition in the host (32). Furthermore, *Lp* induces ROS generation by NADPH oxidase (33), and protects from damaging agents (34). Remarkably, many host responses to *Lactobacilli* are conserved across large evolutionary distances, as *Lactobacillus* strains also coordinate nutrient acquisition (32), ROS generation (33), growth and gut defenses in the mouse (32, 34). These observations position the fly as a valuable model to examine developmental and homeostatic contributions of *Lactobacilli* to animal health (35).

Our interest in *Lp* arose from previous data that elimination of the *Drosophila* microbiome slows ISC turnover, and extends adult longevity (10, 30, 36, 37). These observations led us to ask if symbiotic bacteria revert the germ free (GF)-mediated extension of fly lifespan by accelerating the division of ISCs. To test this hypothesis, we examined the effects of common fly symbionts (38) on GF host longevity. Of all species tested, we found that *Lp* recapitulated the microbiome-mediated truncation of GF adult lifespan. However, counter to our initial expectation, we did not find that *Lp* accelerated ISC division rates. Instead, we found that mono-association of adult flies with *Lp* led to a loss of ISCs, a block to ISC renewal, and a gradual deterioration of epithelial integrity upon aging. Combined, our data show that long-term association of adult ex-GF *Drosophila* with *Lp* destabilizes the intestine, and shortens host longevity.

## RESULTS

### *Lactobacillus plantarum* outcompetes *Lactobacillus brevis* for association with adult *Drosophila*

Our lab strains of *Drosophila* predominantly associate with *Lactobacillus plantarum* (*Lp*), *Lactobacillus brevis* (*Lb*), and *Acetobacter pasteurianus* (*Ap*) (38). Of those strains, *Lactobacilli*, particularly *Lp*, are the dominant symbionts, typically accounting for >75% of all bacterial OTUs in flies that we raise on standard cornmeal medium. As fly symbionts regularly cycle from the intestine to the food (40, 41), we conducted a longitudinal study of the association of *Lp* and *Lb* with cultures of wild-type *Drosophila*. For this work, we fed freshly emerged adult flies an antibiotic cocktail to eliminate the endogenous bacterial microbiome (36, 42). We then fed antibiotic-treated adult flies equal doses of *Lp* or *Lb* for sixteen hours, transferred flies to fresh food, and determined bacterial titers in the intestine and food at regular intervals thereafter (Figure 1A).

**Figure 1.**
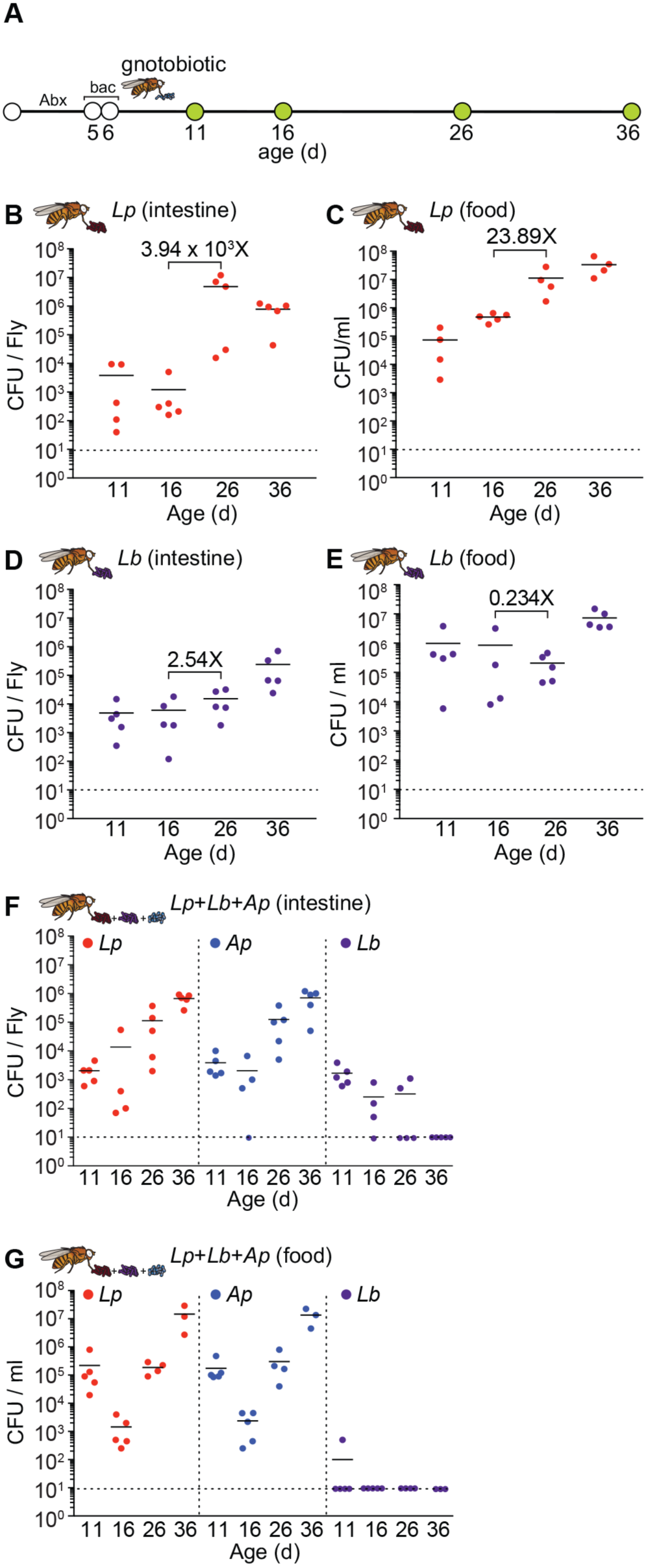
*Lp* outcompetes *Lb* in the adult gut. **(A)** Schematic representation of the experimental timeline and generation of gnotobiotic adult flies. Colony forming units per fly of *Lp* and *Lb* in the intestines **(B, D)** and on the food **(C, E)** of *Lp* mono-associated and *Lb* mono-associated adult flies respectively at days 11 and 16, 26, and 36 of age. Numbers in black indicate fold change in the mean compared between the indicated time points. Colony forming units per fly of *Lp, Lb*, and *Ap* in the **(F)** intestines and on the **(E)** food of *Lp, Lb*, and *Ap* poly-associated adult flies.

We typically found less that 1X10^4^ CFU per fly gut five days after inoculation with either *Lp* or *Lb* (Figure 1B and 1D). In both cases, intestinal bacterial loads increased over time. However, the effect was more pronounced for *Lp* than *Lb.* We detected a mean 4X10^4^ fold increase in numbers of *Lp* associated with the fly gut between days 16 and 26, rising to approximately 1X10^7^ CFU per fly gut by day 26. In contrast, we only observed a 2.5-fold increase in *Lb* gut association over the same time, yielding less than 1X10^5^ CFU per fly gut. Likewise, we found that *Lp* load steadily increased in the food over time (Figure 1C), while the association of *Lb* with food remained relatively constant (Figure 1E). These observations suggest that *Lp* has a growth advantage over *Lb* when co-cultured on fly food with adult *Drosophila*. To determine if *Lp* outcompetes *Lb* for association with *Drosophila*, we fed germ-free adult flies a 1:1:1 mixed culture of *Lp, Lb* and *Ap*, and monitored bacterial association rates over time. We added *Ap* to the culture in this experiment to more accurately represent the microbiome of our conventional lab flies. Of this defined bacterial community, we found that *Lp* and *Ap* populated the fly intestine (Figure 1F) and food (Figure 1G) with near equal efficiency. In both cases, the microbial load associated with the gut or food increased over time, typically reaching approximately 1X10^6^ CFU per intestine 36 days after inoculation. Intestinal association by *Lp* was an order of magnitude higher in mono-associated flies (Figure 1B) than in poly-associated flies (Fig. 1F), suggesting that *Ap* partially limits host association with *Lp*. In contrast to *Lp* and *Ap*, we found that *Lb* gradually disappeared from the food and the intestines of poly-associated adult flies over time (Figure 1F, G). By 36 days, we repeatedly failed to detect *Lb* in the intestine or food. Combined, these observations suggest that the *Lp* and *Ap* strains used in this study are more effective at forming stable, long-term associations with *Drosophila* than the *Lb* strain, and may explain the predominance of *Lp* and *Ap* in fly cultures.

### Mono-association with *Lactobacillus plantarum* shortens adult longevity relative to germ-free counterparts

We and others find that conventionally reared (CR) flies have a shorter lifespan that GF counterparts (30, 36, 43). As *Lp* and *Ap*, readily establish long-term associations with adults (Figure 1), we determined if either species affects the lifespan of adult flies. To address this question, we measured the longevity of adult flies that we associated exclusively with *Lp*, or *Ap*. There are two established methods for the association of *Drosophila* with defined bacterial communities. In one case, embryos are incubated in a bleach solution to remove all associated microbes, and then inoculated with bacteria at the appropriate developmental stage. Alternatively, CR adults are fed a medium treated with a cocktail of antibiotics to eliminate gut bacteria. Antibiotic-treated adults are then fed bacteria to establish a gnotobiotic population (e.g. Figure 1A). Although methodologically distinct, the differences in intestinal transcription between adults derived from bleached embryos and adults treated with antibiotics has not yet been compared. To address this question, we compared microarray data on microbe-dependent gene expression in the intestines of Oregon R and Canton S flies derived from bleached embryos (11), and in the intestines of *w*; *esgGAL4, GAL80^ts^, UAS-GFP* (*esg^ts^*) antibiotic-treated adults (44). The *esg^ts^* genotype is a variant of the *Drosophila* TARGET system (45), and is commonly used for temperature-dependent expression of UAS-bearing transgenes in GFP-marked progenitor cells (46, 47). We used PANTHER to identify GO terms that were significantly enriched in CR intestines relative to GF intestines for all three fly lines. In this comparison, we did not see clear distinctions between the effects of bleach and antibiotics on transcriptional outputs from the gut (Figure 2A, B). In each case, removal of the microbiome altered the expression of immune response genes (Figure 2B), a result that matches earlier data linking gut bacteria and intestinal immunity (11). Of the remaining microbe-responsive GO terms, we noticed a more pronounced similarity between bleached Oregon R flies, and antibiotic-treated *esg^ts^* flies. Of the 21 processes affected by bleach treatment of Oregon R cultures, 10 were similarily affected by antibiotic treatment of *esg^ts^* flies (Figure 2A, B). In contrast, removal of the microbiome with bleach had a mild effect on gut transcription in Canton S flies. In this case, bleach only affected five GO terms, three of which are unique to Canton S (Figure 2A, B). These results suggest that host genetic background is an important determinant of physiological responses to the elimination of commensal microbes in *Drosophila*.

**Figure 2.**
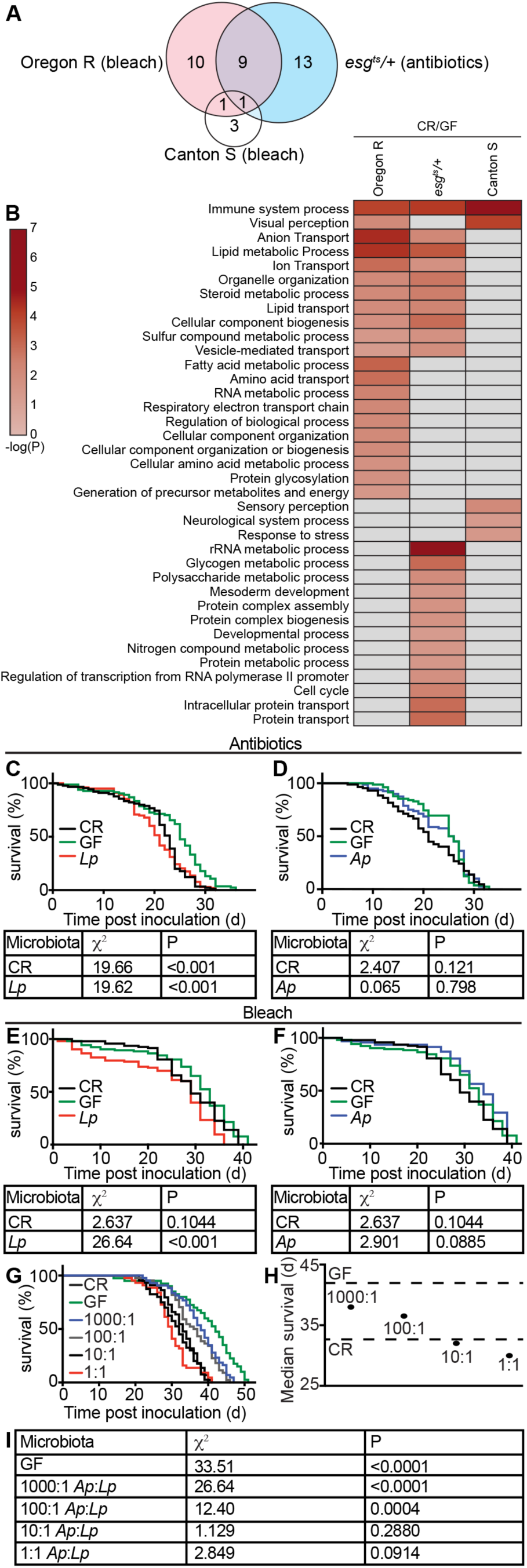
Mono-association with symbiotic *Lp* reduces GF adult fly lifespan. **(A, B)**. Microbe-dependent gene expression microarray data from the intestine of Oregon R and Canton S flies from bleached embryos (11), and in the intestines of *esg^ts^* antibiotic treated adults (63). **(C)** Survival curve of CR, GF, and *Lp* mono-associated adult flies from GF adults generated with antibiotics. **(D)** Survival curve of CR, GF, and *Ap* mono-associated adult flies from GF adults generated with antibiotics. **(E)** Survival curve of CR, GF, and *Lp* mono-associated adult flies generated from bleached embryos. **(F)** Survival curve of CR, GF, and *Ap* mono-associated adult flies generated from bleached embryos. **(G)** Survival curves of CR, GF, and *Ap/Lp* flies co-associated at indicated ratios. For each graph **(C-G)**, the y-axis represents percent survival and the x-axis represents time pot bacterial inoculation. All χ^2^ and p values are relative to GF flies. Tables are results of Long-rank (Mantel-Cox) test for panel data. **(H)** Median survival of data represented in panel **(G). (I)** Comparisons of survival data for the indicated treatment groups relative to CR flies.

To determine if the method of bacterial elimination influences host responses to re-colonization with defined commensals, we prepared GF adults from bleached eggs, or from CR adults raised on antibiotic-treated food. We associated the respective GF adult flies with *Ap* or *Lp*, and measured their lifespans relative to CR counterparts. Irrespective of the means used to generate GF flies, we found that *Lp* significantly shortened the lifespan of adult *Drosophila* (Figure 2C, E). These observations match a recent report that GF adults outlive flies mono-associated with the WJL strain of *Lp* (48) and indicate that mono-association of adults with *Lp* reverts the extended lifespan noted in GF flies. In contrast, mono-association of adult *Drosophila* with *Ap* had no effect on adult lifespan regardless of the method used to generate GF flies (Figure 2D, F). As *Ap* attenuates gut colonization by *Lp* (Fig. 1F) and *Ap* does not affect adult lifespan, we tested if *Ap* attenuates the impacts of *Lp* on GF lifespan extension. For these assays, we measured the lifespans of GF adults that we cultured with different ratios of *Ap* and *Lp*. Here, we observed a clear relationship between *Ap*:*Lp* input ratios and adult lifespan - the greater the ratio of *Ap* to *Lp*, the longer the lifespan of co-associated flies (Figure 2G-I). Consistent with these findings, a recent study also reported that GF flies outlive *Lp*-associated flies, and that addition of *Ap* limits this effect (49). Together, these data argue that mono-association with *Lp* reverts the lifespan extension observed in GF flies.

### *Lp* does not activate proliferative responses in the host intestine

In *Drosophila*, symbiotic bacteria provide mitogenic cues that accelerate the growth and aging of intestinal tissues (10), a factor that may truncate host longevity. This prompted us to test if *Lp* activates ISC division. Initially, we quantified expression of the EGF ligand *spitz* and the *spitz*-activating endopeptidase *rhomboid* in dissected intestines. We selected the EGF pathway for this study, as EGF activates ISC proliferation in response to symbiotic bacteria (10), and damage to the intestinal epithelium (50). Consistent with a relationship between gut bacteria and ISC proliferation, we detected significantly higher levels of *spitz* (Figure 3B), and *rhomboid* (Figure 3C) in CR flies than in GF flies. In contrast, we did not observe expression of EGF pathway activators in the intestines of flies associated with *Lp* (Figure 3B, C). Instead, we found that *spi* was expressed at significantly lower levels in the midguts of *Lp* mono-associated flies than in GF flies by day fifteen (Figure 3D). These data suggest that mono-colonization of the adult intestine with *Lp* does not activate EGF-dependent proliferative responses in the host intestine.

**Figure 3.**
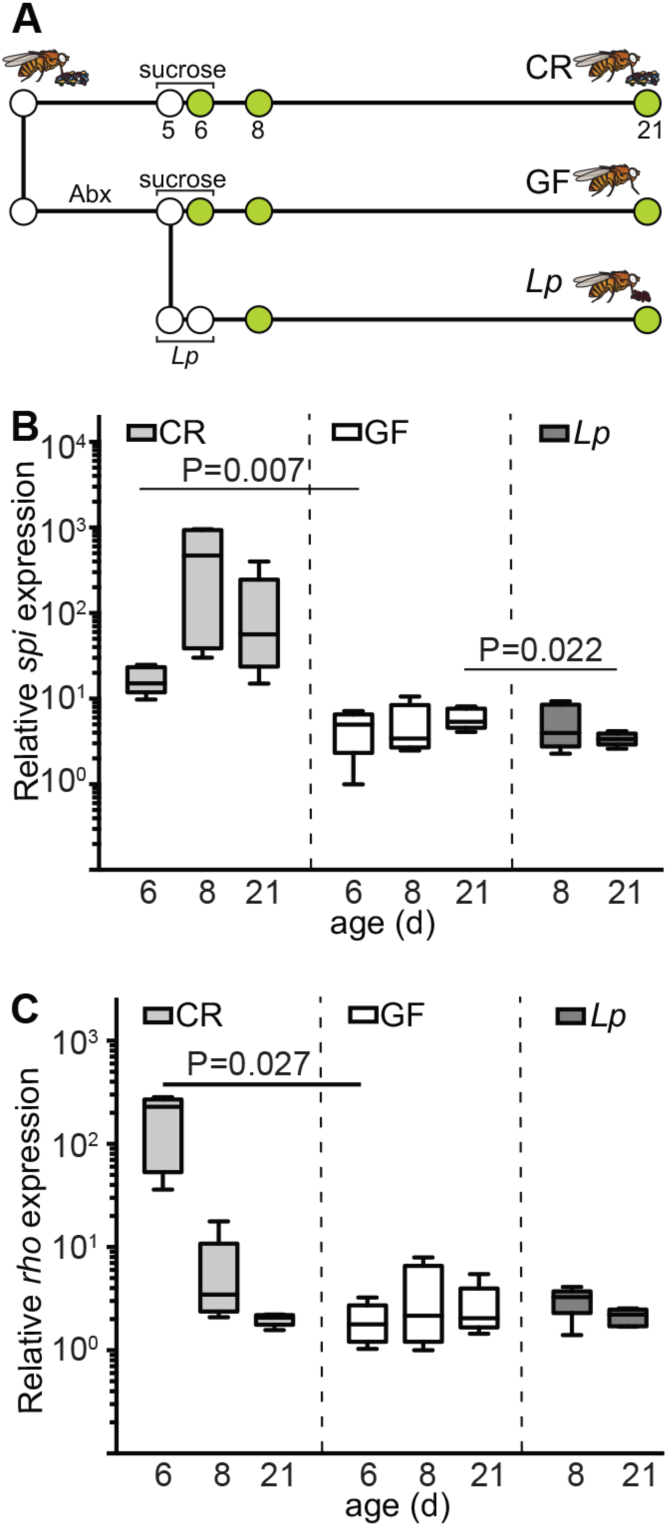
*Lp* does not trigger a proliferative response in adult fly intestines. **(A)** Schematic representation of gnotobiotic fly generation and experimental timeline. Quantitative real-time PCR analysis of expression of **(B)** growth factor *spitz* **(C)** and endopeptidase rhomboid from the dissected guts of adult CR, GF, and *Lp* gnotobiotic flies. Each time point represents five independent measurements.

### Impaired epithelial renewal in *Lp* mono-associated flies

To more accurately determine the effects of *Lp* on ISC proliferation, we used the MARCM clonal marking method to assess stem cell proliferation in the intestines of CR, GF, and *Lp*-associated flies. MARCM labels all progeny of an ISC division with GFP (51). As a result, clone number and size provides a simple proxy for total divisions in the midgut. We looked at ISC division in CR flies, GF flies and flies that we associated with *Lp*. In each case, we counted the total number of mitotic clones per posterior midgut, and the number of cells per clone. As expected, we noticed more mitotic activity in the intestines of CR flies than GF flies. CR flies had significantly more mitotic clones than GF counterparts (Figure 4A, B, D), and CR clones contained significantly more cells than GF clones (Figure 4E). In contrast to CR flies, mono-association with *Lp* failed to initiate proliferative responses in the host (Figure 4C). In fact, the midgut contained significantly fewer clones that CR flies, or GF flies (Figure 4D), and the clones that we observed in *Lp*-associated flies invariably had fewer cells than age-matched clones in CR flies (Figure 4E). These results, in conjunction with our quantitative measurements of host gene expression (Figure 3), demonstrate a near complete absence of epithelial renewal in intestines associated exclusively with *Lp*.

**Figure 4.**
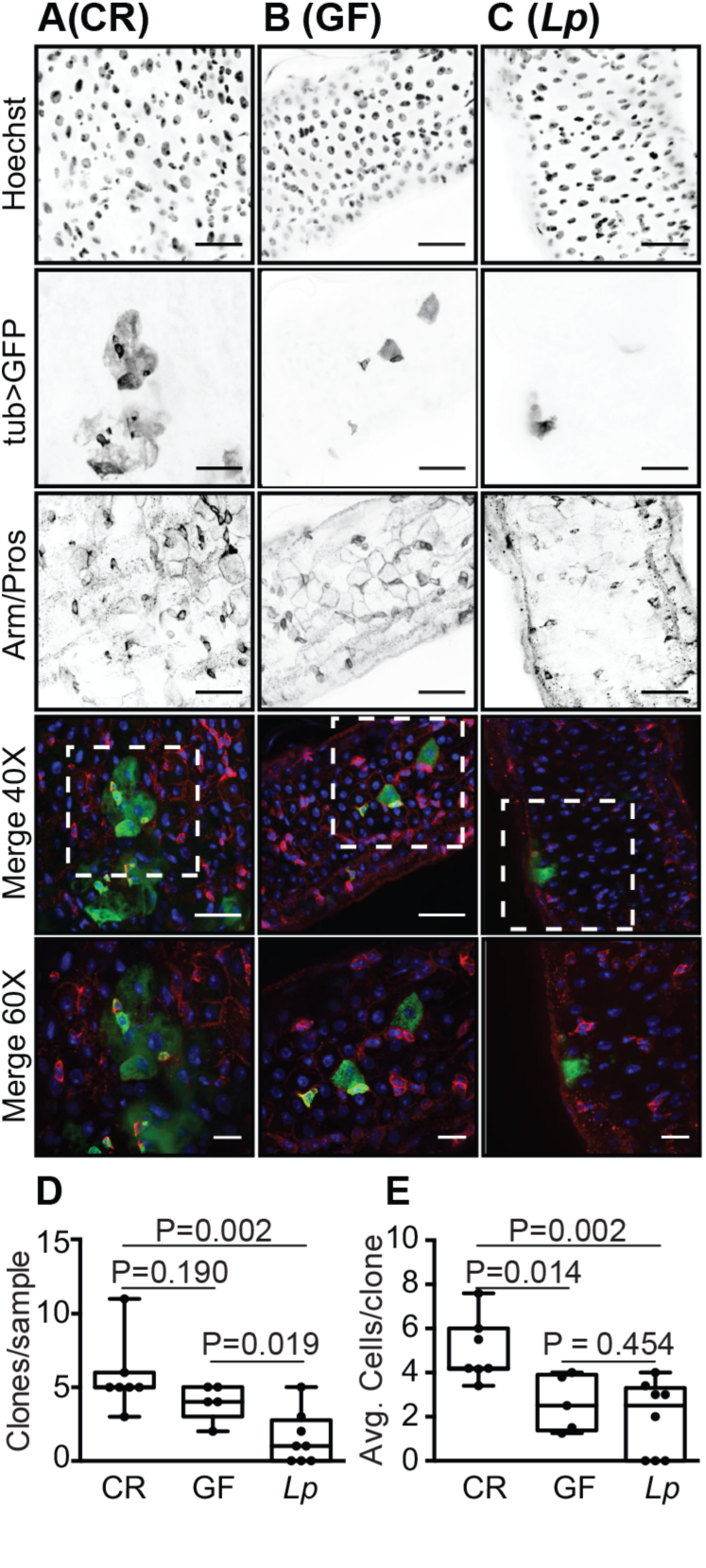
Hindered intestinal regeneration in the guts of *Lp* mono-associated flies. GFP-positive MARCM clones from the posterior midgut of **(A)** CR, **(B)** GF, and **(C)** *Lp* mono-associated flies at day 15-post inoculation. Guts were stained with Hoechst and anti-Arm/Pros antibodies as indicated. Hoechst (blue) and GFP (green) and Arm/Pros (red) were merged in the fourth (40x) and fifth (60x) rows. 40X Scale bars are 25 μm. 60X Scale bars are 5 μm. (D) Quantification of **(D)** clones per sample and **(E)** cells per clone in CR, GF and *Lp* flies.

### *Lp* mono-associated flies lack intestinal progenitors

Given the absence of ISC proliferation, we used immunofluorescence to determine if prolonged mono-association with *Lp* affected the cellular organization of posterior midguts. To measure the influence of *Lp* on midgut morphology, we visualized the posterior midguts of CR, GF, and *Lp* mono-associated *esg^ts^* flies that we raised for two or fifteen days. We used GFP fluorescence, and anti-Armadillo and anti-Prospero immunofluorescence to visualize progenitor cells, cell borders, and enteroendocrince cells, respectively. We did not observe differences between the different treatment groups at the early time point (Fig. 5A-C). In each case, midguts displayed the hallmarks of young intestines - evenly spaced nuclei, regular arrangements of GFP positive progenitors, and neatly organized cell boundaries. As expected, fifteen day old CR midguts showed signs of age-dependent dysplasia (Fig. 5D). We no longer observed regular spacing between individual nuclei, Prospero and Armadillo stains revealed a disorganized epithelium, and the population of GFP-positive progenitors had expanded relative to two day old guts. Consistent with bacterial contributions to the aging of the host intestine, we did not see a similar degree of dysplasia in GF flies. GF flies had regularly spaced nuclei, an organized epithelium, and fewer GFP-positive progenitor cells (Fig. 5E). We also saw minimal signs of dysplasia in the intestines of flies that we associated with *Lp* for fifteen days. In this case, we observed regularly spaced nuclei, defined cell borders, and an even distribution of enteroendocrine cells at day 15 (Fig. 5F). However, we noticed that *Lp*-associated guts had approximately half the progenitor number of GF guts, and significantly fewer than CR guts (Fig. 5G). These data suggest a detrimental impact of *Lp* mono-association on the pool of progenitor cells in adult flies.

**Figure 5.**
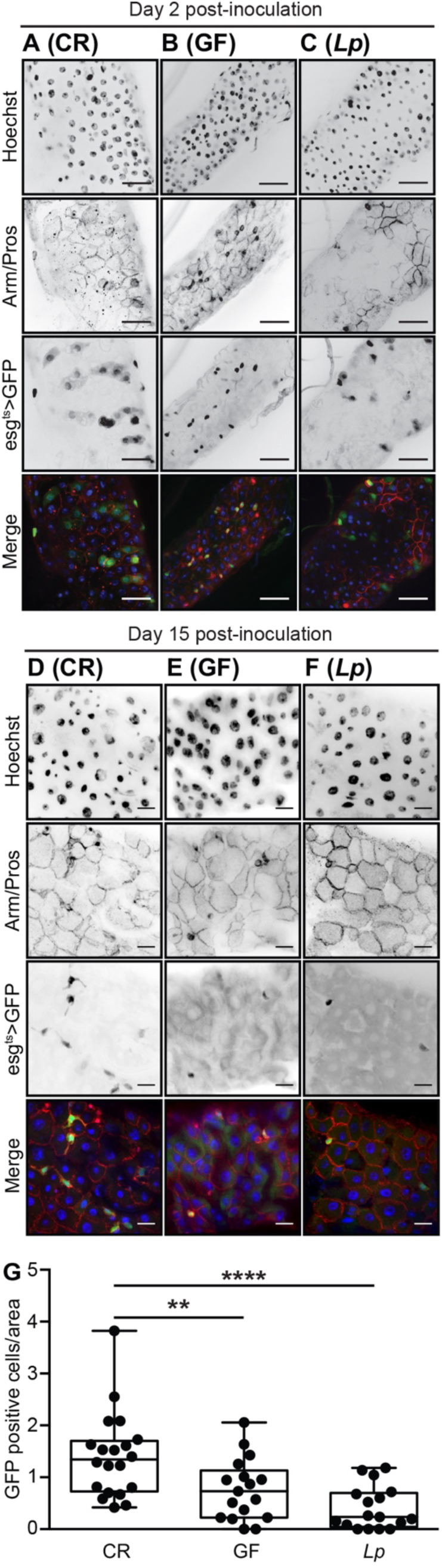
*Lp* mono-associated guts have low numbers of intestinal progenitor cell. Immunofluorescence of posterior midguts of **(A)** CR, **(B)** GF, and **(C)** *Lp* mono-associated associated flies at day 2 post inoculation. Scale bars are 25 μm. Immunofluorescence of posterior midguts of **(D)** CR, **(E)** GF, and **(F)** *Lp* mono-associated associated flies at day 15 post inoculation. Scale bars are 10 μm. Guts were stained with Hoechst and anti-Arm/Pros antibodies as indicated. Progenitor cells were visualized with GFP as indicated. Hoechst (blue), GFP (green), and anti-Arm/Pros (red) were merged in the fourth and eighth row. **(G)** Quantification of progenitor numbers per unit surface area.

### *Lp* disrupts posterior midgut ultrastructure

As mono-association with *Lp* results in a loss of intestinal progenitors, and a failure in epithelial renewal, we used transmission electron microscopy to directly examine the effects of fifteen days mono-association with *Lp* on posterior midgut ultrastructure. As controls, we visualized the posterior midguts of age-matched CR and GF flies. CR midguts had the anticipated sheath of visceral muscle that surrounds basal progenitor cells (Fig. 6A, B). Progenitors, in turn, support the generation of columnar epithelial cells that face the intestinal lumen (Fig. 6B, C). In many ways, GF flies mirrored CR flies, with an organized visceral musculature (Fig. 6D), basal progenitor cells (Fig. 6D, E) and an intact brush border (Fig. 6F). Upon examination of midguts associated with *Lp*, we were struck by substantial alterations to intestinal morphology. The epithelium contained an undulating population of cells (Fig. 6G, H) with large vacuoles (Fig. 6G-I, arrowheads) and poorly-discernible nuclei (Fig. 6G). We also noticed severe alterations to the morphology of presumptive progenitor cells. In place of the small, densely stained progenitors intimately associated with the visceral muscle of CR or GF flies, mono-association with *Lp* resulted in the appearance of misshapen cells that did not associate properly with the muscle, had large, lightly stained nuclei, and numerous cytosolic vacuoles (Fig. 6G, H). These findings show that mono-colonization of a GF adult midgut with *Lp* causes an intestinal phenotype that is characterized by thinning of the epithelium, formation of large cytosolic vacuoles, and a loss of progenitor cells. In summary, mono-association of adult *Drosophila* with *Lp* results in an intestinal phenotype that is distinct from CR or GF flies. *Lp* forms a stable, association with GF *Drosophila* that impairs epithelial renewal programs, depletes progenitor cell populations, and ultimately shortens host longevity.

**Figure 6.**
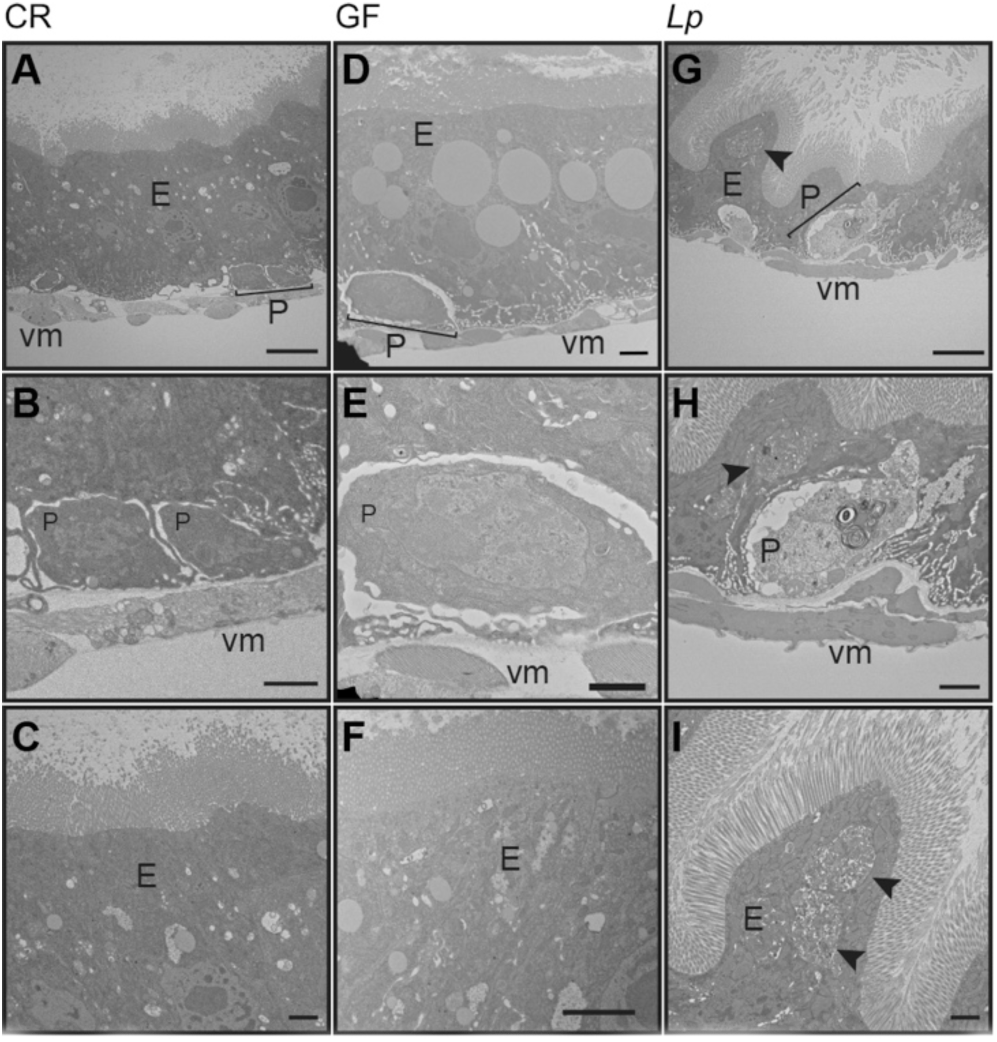
Lp disrupts posterior midgut ultrastructure. Transmission electron microscopy of **(A-C)** CR, **(D-F)** GF, and **(G-I)** *L*p mono-associated posterior midguts 15 days after inoculation. Epithelium (E), progenitors (P), and visceral muscle (VM) are labeled. **(A, E, G)** 1200x direct magnification. Scale bars are 5 μm. **(B, E, H)** 3000x direct magnification. Scale bars are 1μm. **(C, F, I)** 3500x direct magnification. Scale bars are 1μm.

## DISCUSSION

Gut bacteria activate homeostatic division programs in basal intestinal stem cells that are critical for the maintenance of an effective epithelial barrier (10). Failure to regulate stem cell division exposes the host to microbial invasion, and potentiates the development of chronic inflammatory illnesses (52, 53). In this report, we used the *Drosophila* model to examine the effects of symbiotic bacteria on the long-term maintenance of the epithelial renewal program. A recent report showed that mono-association with *Lp* is not deleterious for adult fly fitness, and has benefits for males raised under conditions of nutrient scarcity (48). In this study, we asked if *Lp* affects ISC division independent of effects on lifespan, as we showed previously that manipulation of ISC division rates does not have substantial effects on lifespan (36). To our knowledge, our data are the first to show an impact of mono-association with a common fly symbiont on the maintenance or proliferation of ISCs. We found that mono-association with *Lp* depletes ISC pools, blocks epithelial renewal, and damages the intestinal epithelium. A previous study showed that *gluconobacter morbifer* causes disease in adult *Drosophila* if allowed to expand within the host (54). However, *G. morbifer* is a comparatively rare symbiont of *Drosophila*, and disease onset requires impaired immunity within the host. In contrast, this report identifies a novel intestinal phenotype associated with mono-association of a common fly symbiont with a GF host. We believe these findings represent a valuable model to define the mechanistic basis for stem cell damage by a symbiont.

At present, we do not know how mono-association with *Lp* causes an intestinal pathology within the host. It is possible that this phenotype arises from collateral damage through chronic expression of toxic immune effector molecules such as reactive oxygen species. This hypothesis is supported by the observation that *Lp* activates NADPH-oxidase in the *Drosophila* intestine. Alternatively, errant intestinal immune responses through the Immune Deficiency (IMD) pathway may account for *Lp*-dependent pathologies. In this context, we consider it important to consider that several transcriptional studies demonstrated that a relatively small fraction of IMD-responsive transcripts are easily categorized as bacteriostatic or immune modulatory (55). In fact, it seems that intestinal IMD activity primarily modifies metabolic gene expression (11, 44, 56). As intestinal microbes are known to control nutrition and metabolism in their *Drosophila* host (6), we consider it possible that the *Lp*-dependent pathologies described in this study reflect an underlying imbalance in IMD-dependent regulation of host metabolism. Consistent with possible links between *Lp*, IMD and host metabolism, it is noteworthy that a recent study established a direct link between *Lp* and the IMD-dependent expression of intestinal peptidases (31). Our data show that intestinal colonization by *Lp* is much greater in mono-associated flies than in poly-associated flies. We speculate that the elevated levels of *Lp*, combined with the absence of additional symbionts, alters metabolic responses in the host, leading to impaired intestinal function. This hypothesis includes the possibility that *Lp* directly affects host diet, as proposed for other *Drosophila*-associated microbes (57, 58). We are currently testing this hypothesis in flies with modified intestinal IMD activity.

Our work was initially inspired by reports from our group and others that GF adults outlive CR flies (36, 59). However, other studies reported variable impacts of the effects of microbiome removal on adult lifespan (60, 61). We believe that the differences between the individual reports reflects the intricate nature of interactions within a host-microbe-environment triad. For example, research groups typically raise their flies on an incompletely defined diet that exerts underappreciated influences on the metabolic outputs of intestinal bacteria, and the transcriptional outputs of the host. We believe that a complete evaluation of the relationship between microbes and their hosts requires consideration of environmental inputs such as diet. Likewise, it is important to consider genotypic inputs of the symbiont strain and the fly host. For example, the beneficial contributions of *Lp* to mouse and larval nutrition display strain-specific effects (9, 32), suggesting variable effects of individual *Lp* strains on host phenotypes. In addition, most research groups study lab-raised strains of experimental models, a situation that allows unexplored genetic drift within the host to influence microbiome composition and phenotype.

In summary, this report uncovers long-term negative effects of *Lactobacillus plantarum* on the maintenance and growth of the intestinal stem cell pool. Given the experimental accessibility of *Drosophila* and *Lactobacilli*, we believe these findings represent a valuable tool for the definition of the mechanisms by which individual symbionts influence stem cell homeostasis.

## MATERIALS AND METHODS

### Bacterial Strains

All *Drosophila* symbiotic bacteria strains used were isolated from wild-type lab flies from the Foley lab at the University of Alberta and are as follows: *Lactobacillus plantarum* KP (DDBJ/EMBL/GenBank chromosome 1 accession CP013749 (https://www.ncbi.nlm.nih.gov/nuccore/CP013749) and plasmids 1-3 for accession numbers CP013750 (https://www.ncbi.nlm.nih.gov/nuccore/CP013750), CP013751 (https://www.ncbi.nlm.nih.gov/nuccore/CP013751), and CP013752 (https://www.ncbi.nlm.nih.gov/nuccore/CP013752), respectively), *Lactobacillus brevis* EF (DDBJ/EMBL/GeneBank accession LPXV00000000 (https://www.ncbi.nlm.nih.gov/nuccore/LPXV00000000)), and *Acetobacter pasteurianus* AD (DDBJ/EMBL/GeneBank accession LPWU00000000 (https://www.ncbi.nlm.nih.gov/nuccore/LPWU00000000) and are described in (38). *Lactobacillus* strains were grown in MRS broth (Sigma Lot: BCBS2861V) at 29°C and *Acetobacter pasteurianus* was grown in Mannitol broth (2.5% n-mannitol. 0.5% yeast extract, 0.3% peptone) 29°C with shaking.

### Colony forming units per fly

At indicated time points 25 flies were collected from an indicated group and placed into successive solutions of 20% bleach, distilled water, 70% ethanol, distilled water to surface sterilize and rinse flies respectively. These 25 flies were then randomly divided into groups of 5 and mechanically homogenized in MRS broth. Fly homogenate was then diluted in serial dilutions in a 96 well plate and 10ul spots were then plated on either MRS agar (to select for *Lactobacillius*) GYC agar (to select for *Acetobacter*). Plates were incubated for 2 days at 29°C and the number of colonies per bacterial species were counted. Bacterial species were distinguished by colony morphology

### Fly husbandry

All experiments were performed with virgin female flies. *w^1118^* flies were used as wild-type strain and used in all experiments unless otherwise mentioned. The *w, esg-GAL4, tubGAL80[ts]]* flies have previously been described (46, 47). Mitotic clones were generated with flies of the genotype *y,w, hs-flp, UAS-mCD8GFP; neoFRT(40A)/neoFRT(40A),tubGAL80; tubGAL4/+.* Flies were raised on standard corn meal medium (Nutri-Fly Bloomington Formulation, Genesse Scientific) at 29° C. Germ-Free flies were generated by antibiotic treatment were made germ free by raising adult flies on autoclaved standard media supplemented with an antibiotic solution (100 μg/ml Ampicillin, 100 μg/ml Metronidazole, 40 μg/ml Vancomycin dissolved in 50% ethanol and 100 μg/ml Neomycin dissolved in water) to eliminate the microbiome from adult flies (54). CR flies were raised on autoclaved standard cornmeal medium. To obtain axenic fly stocks from embryo, embryos were laid on apple juice plates over a 16-h period and then collected. All the following steps were performed in a sterile hood. Embryos were rinsed from the plate with sterile PBS. Embryos were placed in 10% sodium hypochlorite solution for 2.5 min and then placed into a fresh 10% sodium hypochlorite solution for 2.5 min and then washed with 70% EtOH for 1 min. Embryos were then rinsed 3 times with sterile water and placed onto sterile food and maintained at 25°C in a sterilized incubator in a sterile hood. Axenic flies were generated in parallel with conventionally reared counterparts who were placed in water at all steps. For adult fly longevity studies N of 100 (Fig. 2) flies were raised in vials with 20 flies per vial. Flies were transferred to fresh food every 2-3 days.

### Generation of gnotobiotic *Drosophila*

Virgin females were raised on antibiotic supplemented medium for 5 days at 29° C. On day 5 of antibiotic treatment, a fly from each group to be mono/co/ploy-associated was homogenized in MRS broth and plated on MRS and GYC agar plates to ensure eradication of pre-existing microbes. Flies were starved in sterile empty vials for 2 hours prior to bacterial association. For mono-associations, the OD_600_ of bacteria liquid cultures was measured and then the culture was spun down and re-suspended in in 5% sucrose in PBS to final OD_600_ of 50. For axenic embryos, virgin female flies were collected for 2-3 days and then associated with the same protocol as antibiotic GF treated flies. For co-associations, bacterial cultures of *Ap* and *Lp* were prepared to an OD_600_ of 50 in 5% sucrose in PBS as described above. The bacterial cultures were then mixed at ratios of 1000:1, 100:1, 10:1, and 1:1 *Ap* to *Lp.* For poly-associations, bacterial cultures of *Ap, Lb*, and *Lp* were prepared to an OD_600_ of 50 in 5% sucrose in PBS as described above. The bacterial cultures were then mixed at a 1:1:1 ratio. For all bacterial associations, twenty-two flies/vial were associated with 1ml of bacterial suspension on autoclaved cotton plugs. Flies were fed bacteria sucrose mixture for 16 hours at 29°C and then kept on autoclaved food for the remainder of the study. CR and GF flies were given mock associations of 1ml of 5% sucrose in PBS for 16 hours at 29°C. To ensure mono-association or GF conditions, respective flies were homogenized in MRS broth and plated on MRS or GYC agar plates periodically throughout the study.

### Immunofluorescence

Flies were washed with 95% ethanol and dissected in PBS to isolate adult intestines. Guts were fixed for 20 minutes at room temperature in 5% formaldehyde in PBS. Guts were rinsed in PBS for 20 minutes at room temperature and blocked overnight in PBSTBN (PBS, 0.05% Tween 20, 5% BSA, and 1% goat serum) at 4°C. Guts were stained overnight at 4°C in PBSTBN with appropriate antibodies, washed with PBSTB (PBS, 0.05% Tween 20, and 5% BSA) and stained for 1 h at room temperature in PBSTBN with Hoechst 33258 (1:500) and the appropriate secondary antibody (goat anti-mouse Alexa Fluor 568 or goat anti-mouse Alexa Fluor 647) guts were washed with PBSTB and rinsed with PBS prior to visualization. The primary antibodies used in this study were as follows: mouse anti-Armadillo (1:100; Developmental Studies Hybridoma Bank N2 7A1), mouse anti-Prospero (1:100; Developmental Studies Hybridoma Bank MR1A). Guts were mounted on slides in Fluoromount (Sigma-Aldrich F4680), and the posterior midgut was visualized with a spinning disk confocal microscope (Quorum WaveFX, Quorum Technologies Inc.). Images were collected as z-slices and processed with Fiji software to generate a single *Z*-stacked image.

### qPCR

Realtime PCR was performed on the dissected guts of adult *Drosophila*. The quantitative PCR protocol and primers used in this study have been described previously (62).

### Transmission Electron Microscopy

Flies were washed with 95% ethanol and dissected into PBS. Posterior midguts were immediately excised and placed into fixative (3% paraformaldehyde + 3% glutaraldehyde). Fixation preparation, contrasting sectioning, sectioning, and visualization were performed at the Faculty of Medicine and Dentistry Imaging Core at the University of Alberta. Midgut sections were visualized with Hitachi H-7650 transmission electron microscope at 60Kv in high contrast mode.

## ACKNOWLEDGMENTS

We are grateful to Dr. François Leulier for comments on the manuscript, and to Dr. Nichole Broderick for the Oregon R and Canton S microarray data presented in figure 2. Transgenic flies were provided by Dr. Bruno Lemaitre. We are also grateful to Kristina Petkau for her aid in the generation of axenic embryos. We acknowledge the microscopy support from Dr. Stephen Ogg, Woo Jung Cho, and the Faculty of Medicine and Dentistry core imaging service, the Cell Imaging Centre, University of Alberta.

## AUTHOR CONTRIBUTIONS

D.F and E.F. conceived and designed experiments; D.F and A.D. performed the experiments; D.F. and E.F performed data analysis and wrote the paper.

## REFRENCES

1. Hacquard S, Garrido-Oter R, González A, Spaepen S, Ackermann G, Lebeis S, McHardy AC, Dangl JL, Knight R, Ley R, Schulze-Lefert P. 2015. Microbiota and Host Nutrition across Plant and Animal Kingdoms. Cell Host Microbe 17:603–616.

2. Hooper L V, Littman DR, Macpherson AJ. 2012. Interactions between the microbiota and the immune system. Science 336:1268–73.

3. Hsiao EY, McBride SW, Hsien S, Sharon G, Hyde ER, McCue T, Codelli JA, Chow J, Reisman SE, Petrosino JF, Patterson PH, Mazmanian SK. 2013. Microbiota Modulate Behavioral and Physiological Abnormalities Associated with Neurodevelopmental Disorders. Cell 155:1451–1463.

4. Kamada N, Seo S-U, Chen GY, Núñez G. 2013. Role of the gut microbiota in immunity and inflammatory disease. Nat Rev Immunol 13:321–335.

5. Kamada N, Chen GY, Inohara N, Núñez G. 2013. Control of pathogens and pathobionts by the gut microbiota. Nat Immunol 14:685–90.

6. Wong AC-N, Dobson AJ, Douglas AE. 2014. Gut microbiota dictates the metabolic response of Drosophila to diet. J Exp Biol 217:1894–901.

7. Blumberg R, Powrie F. 2012. Microbiota, Disease, and Back to Health: A Metastable Journey. Sci Transl Med 4.

8. Carding S, Verbeke K, Vipond DT, Corfe BM, Owen LJ. 2015. Dysbiosis of the gut microbiota in disease. Microb Ecol Health Dis 26:26191.

9. Storelli G, Defaye A, Erkosar B, Hols P, Royet J, Leulier F. 2011. Lactobacillus plantarum Promotes Drosophila Systemic Growth by Modulating Hormonal Signals through TOR-Dependent Nutrient Sensing. Cell Metab 14:403–414.

10. Buchon N, Broderick NA, Chakrabarti S, Lemaitre B. 2009. Invasive and indigenous microbiota impact intestinal stem cell activity through multiple pathways in Drosophila. Genes Dev 23:2333–44.

11. Broderick NA, Buchon N, Lemaitre B. 2014. Microbiota-Induced Changes in Drosophila melanogaster Host Gene Expression and Gut Morphology Microbiota-Induced Changes in Drosophila melanogaster Host Gene. MBio 5:1–13.

12. Matos RC, Leulier F. 2014. Lactobacilli-Host mutualism: “learning on the fly.” Microb Cell Fact 13:S6.

13. Ma D, Storelli G, Mitchell M, Leulier F. Studying Host–microbiota Mutualism in Drosophila: Harnessing the Power of Gnotobiotic Flies.

14. Wong CNA, Ng P, Douglas AE. 2011. Low-diversity bacterial community in the gut of the fruitfly Drosophila melanogaster. Environ Microbiol 13:1889–1900.

15. Wong AC-N, Chaston JM, Douglas AE. 2013. The inconstant gut microbiota of Drosophila species revealed by 16S rRNA gene analysis. ISME J 7:1922–1932.

16. Adair KL, Douglas AE. 2017. Making a microbiome: the many determinants of host-associated microbial community composition. Curr Opin Microbiol 35:23–29.

17. Obadia B, Güvener ZT, Zhang V, Ceja-Navarro JA, Brodie EL, Ja WW, Ludington WB. 2017. Probabilistic Invasion Underlies Natural Gut Microbiome StabilityCurrent Biology.

18. Broderick NA, Lemaitre B. 2012. Gut-associated microbes of Drosophila melanogaster. Gut Microbes 3:307–321.

19. Obadia B, Güvener ZT, Zhang V, Ceja-Navarro JA, Brodie EL, Ja WW, Ludington WB. 2017. Probabilistic Invasion Underlies Natural Gut Microbiome Stability. Curr Biol 27:1999–2006.e8.

20. Lemaitre B, Miguel-Aliaga I. 2013. The Digestive Tract of Drosophila melanogaster. Annu Rev Genet 47:377–404.

21. Buchon N, Osman D, David FPA, Yu Fang H, Boquete J-P, Deplancke B, Lemaitre B. 2013. Morphological and Molecular Characterization of Adult Midgut Compartmentalization in Drosophila. Cell Rep 3:1725–1738.

22. Jiang H, Edgar BA. 2012. Intestinal stem cell function in Drosophila and mice. Curr Opin Genet Dev 22:354–360.

23. Ma D, Storelli G, Mitchell M, Leulier F. Studying Host–microbiota Mutualism in Drosophila: Harnessing the Power of Gnotobiotic Flies.

24. Ohlstein B, Spradling A. 2006. The adult Drosophila posterior midgut is maintained by pluripotent stem cells. Nature 439:470–474.

25. Cheng H, Leblond CP. 1974. Origin, differentiation and renewal of the four main epithelial cell types in the mouse small intestine V. Unitarian theory of the origin of the four epithelial cell types. Am J Anat 141:537–561.

26. Crosnier C, Stamataki D, Lewis J. 2006. Organizing cell renewal in the intestine: stem cells, signals and combinatorial control. Nat Rev Genet 7:349–359.

27. Jin Y, Patel PH, Kohlmaier A, Pavlovic B, Zhang C, Edgar BA. 2017. Intestinal Stem Cell Pool Regulation in Drosophila. Stem Cell Reports 8:1479–1487.

28. Rera M, Clark RI, Walker DW. 2012. Intestinal barrier dysfunction links metabolic and inflammatory markers of aging to death in Drosophila. Proc Natl Acad Sci U S A 109:21528–33.

29. Belkaid Y, Hand TW. 2014. Role of the Microbiota in Immunity and Inflammation. Cell 157:121–141.

30. Clark RI, Salazar A, Yamada R, Fitz-Gibbon S, Morselli M, Alcaraz J, Rana A, Rera M, Pellegrini M, Ja WW, Walker DW. 2015. Distinct Shifts in Microbiota Composition during Drosophila Aging Impair Intestinal Function and Drive Mortality. Cell Rep 12:1656–67.

31. Erkosar B, Storelli G, Mitchell M, Bozonnet L, Bozonnet N, Leulier F. 2015. Pathogen Virulence Impedes Mutualist-Mediated Enhancement of Host Juvenile Growth via Inhibition of Protein Digestion. Cell Host Microbe 18:445–455.

32. Schwarzer M, Makki K, Storelli G, Machuca-Gayet I, Srutkova D, Hermanova P, Martino ME, Balmand S, Hudcovic T, Heddi A, Rieusset J, Kozakova H, Vidal H, Leulier F. 2016. Lactobacillus plantarum strain maintains growth of infant mice during chronic undernutrition. Science 351:854–7.

33. Jones RM, Luo L, Ardita CS, Richardson AN, Kwon YM, Mercante JW, Alam A, Gates CL, Wu H, Swanson PA, Lambeth JD, Denning PW, Neish AS. 2013. Symbiotic lactobacilli stimulate gut epithelial proliferation via Nox-mediated generation of reactive oxygen species. EMBO J 32:3017–28.

34. Jones RM, Desai C, Darby TM, Luo L, Wolfarth AA, Scharer CD, Ardita CS, Reedy AR, Keebaugh ES, Neish AS. 2015. Lactobacilli Modulate Epithelial Cytoprotection through the Nrf2 Pathway. Cell Rep 12:1217–1225.

35. Matos RC, Leulier F. 2014. Lactobacilli-Host mutualism: “learning on the fly.” Microb Cell Fact 13 Suppl 1:S6.

36. Petkau K, Parsons BD, Duggal A, Foley E. 2014. A deregulated intestinal cell cycle program disrupts tissue homeostasis without affecting longevity in Drosophila. J Biol Chem 289:28719–29.

37. Galenza A, Hutchinson J, Campbell SD, Hazes B, Foley E. 2016. Glucose modulates Drosophila longevity and immunity independent of the microbiota. Biol Open 5:165–73.

38. Petkau K, Fast D, Duggal A, Foley E. 2016. Comparative evaluation of the genomes of three common Drosophila-associated bacteria. Biol Open 5:1305–16.

39. Fast D, Duggal A, Foley E. 2016. Lactobacillus plantarum is a pathobiont for adult Drosophila. bioRxiv 49981.

40. Blum JE, Fischer CN, Miles J, Handelsman J. 2013. Frequent replenishment sustains the beneficial microbiome of Drosophila melanogaster. MBio 4:e00860–13.

41. Erkosar B, Leulier F. 2014. Transient adult microbiota, gut homeostasis and longevity: Novel insights from the Drosophila model. FEBS Lett 588:4250–4257.

42. Ryu J-H, Kim S-H, Lee H-Y, Bai JY, Nam Y-D, Bae J-W, Lee DG, Shin SC, Ha E-M, Lee W-J. 2008. Innate immune homeostasis by the homeobox gene caudal and commensal-gut mutualism in Drosophila. Science 319:777–82.

43. Galenza A, Hutchinson J, Campbell SD, Hazes B, Foley E. 2016. Glucose modulates Drosophila longevity and immunity independent of the microbiota. Biol Open 5.

44. Petkau K, Ferguson M, Guntermann S, Foley E. 2017. Constitutive Immune Activity Promotes Tumorigenesis in Drosophila Intestinal Progenitor Cells. Cell Rep 20:1784–1793.

45. McGuire SE, Mao Z, Davis RL. 2004. Spatiotemporal gene expression targeting with the TARGET and gene-switch systems in Drosophila. Sci STKE 2004:pl6.

46. Avadhanula V, Weasner BP, Hardy GG, Kumar JP, Hardy RW. 2009. A Novel System for the Launch of Alphavirus RNA Synthesis Reveals a Role for the Imd Pathway in Arthropod Antiviral Response. PLoS Pathog 5:e1000582.

47. Buchon N, Broderick NA, Kuraishi T, Lemaitre B. 2010. Drosophila EGFR pathway coordinates stem cell proliferation and gut remodeling following infection. BMC Biol 8:152.

48. Téfit MA, Leulier F. 2017. Lactobacillus plantarum favors the early emergence of fit and fertile adult Drosophila upon chronic undernutrition. J Exp Biol 220:900–907.

49. Gould AL, Zhang V, Lamberti L, Jones EW, Obadia B, Gavryushkin A, Carlson JM, Beerenwinkel N, Ludington WB. 2017. High-dimensional microbiome interactions shape host fitness. bioRxiv 232959.

50. Jiang H, Grenley MO, Bravo M-J, Blumhagen RZ, Edgar BA. 2011. EGFR/Ras/MAPK Signaling Mediates Adult Midgut Epithelial Homeostasis and Regeneration in Drosophila. Cell Stem Cell 8:84–95.

51. Wu JS, Luo L. 2007. A protocol for mosaic analysis with a repressible cell marker (MARCM) in Drosophila. Nat Protoc 1:2583–2589.

52. Buchon N, Broderick NA, Chakrabarti S, Lemaitre B. 2009. Invasive and indigenous microbiota impact intestinal stem cell activity through multiple pathways in Drosophila. Genes Dev 23:2333–44.

53. Rera M, Clark RI, Walker DW. 2012. Intestinal barrier dysfunction links metabolic and inflammatory markers of aging to death in Drosophila. Proc Natl Acad Sci U S A 109:21528–33.

54. Ryu J-H, Kim S-H, Lee H-Y, Bai JY, Nam Y-D, Bae J-W, Lee DG, Shin SC, Ha E-M, Lee W-J. 2008. Innate immune homeostasis by the homeobox gene caudal and commensal-gut mutualism in Drosophila. Science 319:777–82.

55. De Gregorio E, Spellman PT, Tzou P, Rubin GM, Lemaitre B. 2002. The Toll and Imd pathways are the major regulators of the immune response in Drosophila. EMBO J 21:2568–79.

56. Combe BE, Defaye A, Bozonnet N, Puthier D, Royet J, Leulier F. 2014. Drosophila Microbiota Modulates Host Metabolic Gene Expression via IMD/NF-κB Signaling. PLoS One 9:e94729.

57. Chaston JM, Newell PD, Douglas AE. 2014. Metagenome-wide association of microbial determinants of host phenotype in Drosophila melanogaster. MBio 5:e01631–14.

58. Yamada R, Deshpande SA, Bruce KD, Mak EM, Ja WW. 2015. Microbes Promote Amino Acid Harvest to Rescue Undernutrition in Drosophila. Cell Rep 10:865–872.

59. Clark RI, Salazar A, Yamada R, Fitz-Gibbon S, Morselli M, Alcaraz J, Rana A, Rera M, Pellegrini M, Ja WW, Walker DW. 2015. Distinct Shifts in Microbiota Composition during Drosophila Aging Impair Intestinal Function and Drive Mortality. Cell Rep 12:1656–67.

60. Brummel T, Ching A, Seroude L, Simon AF, Benzer S. 2004. Drosophila lifespan enhancement by exogenous bacteria. Proc Natl Acad Sci U S A 101:12974–9.

61. Ren C, Webster P, Finkel SE, Tower J. 2007. Increased Internal and External Bacterial Load during Drosophila Aging without Life-Span Trade-Off. Cell Metab 6:144–152.

62. Guntermann S, Foley E. 2011. The protein Dredd is an essential component of the c-Jun N-terminal kinase pathway in the Drosophila immune response. J Biol Chem 286:30284–94.

63. Constitutive Immune Activity Promotes Tumorigenesis in Drosophila Intestinal Progenitor Cells.

